# Identifying molecular features that are associated with biological function of intrinsically disordered protein regions

**DOI:** 10.1101/2020.06.23.167361

**Authors:** Taraneh Zarin, Bob Strome, Gang Peng, Iva Pritišanac, Julie D Forman-Kay, Alan M Moses

## Abstract

Previously, we showed that intrinsically disordered regions in proteins (IDRs) contain multiple sequence-distributed molecular features that are conserved over evolution, despite little sequence similarity that can be detected in alignments (Zarin et al. 2019). Here, we aim to use these molecular features to make specific functional predictions for individual IDRs and identify the molecular features within them that are responsible for specific functions. We find that the predictable functions are diverse, identifying previously known associated molecular features, as well as features that were previously not known to be associated with these functions. We experimentally confirm that elevated isoelectric point and hydrophobicity, features that are positively associated with mitochondrial localization, are necessary for mitochondrial targeting function. Remarkably, increasing isoelectric point in a synthetic IDR restores weak mitochondrial targeting. We believe feature analysis represents a new systematic approach to understand how biological functions of IDRs are specified by their protein sequences.

## Introduction

Intrinsically disordered regions (IDRs), which are protein regions that lack stable tertiary structure, are increasingly appreciated as playing key roles in diverse aspects of cell biology (Forman-Kay and Mittag, 2013). Bioinformatics methods identify thousands of IDRs in eukaryotic proteomes (Dosztányi et al., 2005; Uversky, 2002) and methods have been developed to predict biophysical or structural behavior for specific subsets of IDRs, such as those that fold upon binding (Katuwawala et al., 2019), contain specific N- or C-terminal motifs (e.g., (Chen et al., 2008; Chuang et al., 2012)), or phase separate (e.g. (Vernon et al., 2018), reviewed in (Vernon and Forman-Kay, 2019)). Although there have been recent advances in predicting function from disordered region sequences (reviewed in (Uversky, 2020)), for the vast majority of IDRs it is currently difficult to obtain high-specificity predictions of biological function, or to identify which amino acid residues within them are important for function (Van Der Lee et al., 2014). Progress toward these goals would facilitate designing assays and mutagenesis experiments to understand the function of intrinsically disordered regions, which is especially pressing for those intrinsically disordered regions that are mutated in disease (Vacic and Iakoucheva, 2012).

For structured regions, similarity to homologous protein sequences offers highly specific and diverse predictions of biological function (El-Gebali et al., 2019) and can point to key residues (Ondrechen et al., 2001). However, intrinsically disordered regions often show little sequence similarity detectable in alignments, and therefore alignment-based methods (Davey et al., 2012; Nguyen Ba et al., 2012) reveal only a small part of the functional elements in these regions (Nguyen Ba et al., 2012). Recent studies indicate that despite little amino acid sequence conservation, intrinsically disordered regions (IDRs) contain sequence-distributed molecular features, such as biophysical properties, repeats and short linear motifs, that are likely under natural selection and associated with biological function (Zarin et al., 2017, 2019). This suggests that, using appropriate statistical methods, it should be possible to predict function for individual IDRs based on their amino acid sequences, and to associate specific molecular features with biological functions. If successful, such an approach would allow general predictions of diverse functions for IDRs, and generate testable hypotheses about molecular features that are important for those functions. This contrasts with current bioinformatics approaches that are designed to predict specific IDR subtypes or approaches that predict whole-protein function based on amino acid sequences (Lobley et al., 2007; Radivojac et al., 2013).

Here we describe a first general approach to predict diverse functions for intrinsically disordered regions and to identify molecular features that are important for those functions. Using this approach, we identify molecular features that are associated with functions such as subcellular organization and signaling. This allows us to confirm experimentally that molecular features are necessary for mitochondrial targeting signals, and can be used to establish a mitochondrial targeting signal in a synthetic IDR. We also demonstrate how our approach can predict a role in transcription for the N-terminal IDR in an uncharacterized yeast protein Mfg1, based in part on evolutionary variation in polyglutamine repeats. Based on these examples, our work indicates a path forward for prediction of specific biological functions based on IDR sequences, and experimental characterization of molecular features that are necessary for IDR function.

## Results

### An interpretable, regularized probabilistic model to predict function from IDR features

One of the main challenges of training statistical models to predict IDR function is the lack of systematic data at the IDR level. Functional annotations such as Gene Ontology (The Gene Ontology Consortium, 2019) or deletion phenotypes are at the protein level, but many proteins contain more than one predicted IDR. We therefore devised a statistical model that could combine the contributions of all IDRs in a protein. We assume that IDRs are associated with feature vectors *Z* (for example, the “evolutionary signatures” that summarize the evolution of molecular features (Zarin et al., 2019)) and that functional annotations, *Y*, are binary indicator variables at the protein level (we treat each annotation independently) such that *Y*=1 if the protein is associated with the given annotation, and *Y*=0 if the protein is not associated with the given annotation. If we knew that the *j*-th IDR in the *i*-th protein was responsible for the function, this would amount to a standard classification problem.

Since our goal is to develop a method that can identify the molecular features associated with each function as well as to predict the function of the IDRs, we consider logistic regression, which summarizes the contribution of each feature to the prediction task using a vector of coefficients, *b*, one for each of *m* features. Using standard logistic regression, if the IDR responsible for a function were known we could write the probability of a function given the features

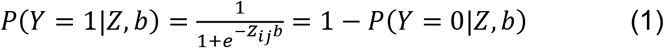

Since, in fact, we do not know which IDR is responsible for the functional annotation *Y*, we treat this as a hidden indicator variable, *X*, and marginalize. The likelihood of the data for *n* proteins is then:

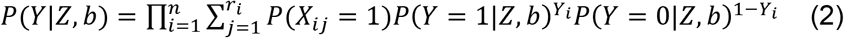

where *r_i_* is the number of disordered regions in the *i*-th protein, and *X_ij_* = 1 if the *j*-th IDR in the *i*-th protein is responsible for the function, *Y*.

To estimate the parameters and infer the hidden variables, we use an E-M (Expectation-Maximization) algorithm (reviewed in (Moses, 2017)). At the E-step, we use Bayes Theorem to compute the expected value of the hidden variable, *X* (see Materials and methods). Because for many functions we have small numbers of positive examples (relative to the number of molecular features), at the M-step we sought to maximize an L1-penalized complete likelihood so that most of the coefficients, *b*, are set to zero (Hastie et al., 2015). We find that the complete likelihood of our model amounts to a weighted logistic regression and therefore, we can use IRLS (Iteratively Re-weighted Least Squares) to do L1 regularized regression at the M-step (see Materials and methods). Thus, the model automatically determines which IDRs are likely to be responsible for each function and which features are most informative for each function.

We refer to this model as FAIDR (for Feature Analysis of Intrinsically Disordered Regions). Given IDR feature vectors, in principle, it can predict protein function (predict *Y*), infer which IDR is responsible for a given function (infer the posterior probability of hidden variable *X*), and determine the molecular features that are important for each function (non-zero components of *b*).

### IDR function can be predicted from protein-level annotations and IDR sequence properties

We trained FAIDR models using protein-level functional annotations from the Saccharomyces Genome Database (SGD) (Cherry et al., 2012) (see Methods). As previously described (Zarin et al., 2019), to summarize the molecular features of IDRs, we used evolutionary signatures, vectors of Z-scores that summarize the deviations of 82 molecular features in disordered regions from the distribution of those features obtained from simulations of IDR evolution. For each molecular feature we calculate the mean and (log) variance across a set of orthologous IDRs, which leads to a total of 164 features describing the evolution of these molecular features in each IDR. Here we wanted to use both the evolutionarily conserved and diverged parts of the disordered region to form the evolutionary signature. Therefore, in contrast to our previous work (Zarin et al., 2019) we did not constrain the simulations to preserve short conserved motifs (see Materials and Methods for details). The evolutionary signatures that we calculated for the yeast proteome are available for download and visualization at www.moseslab.csb.utoronto.ca/idr/.

To demonstrate the power of FAIDR at predicting IDR-level function, we used two systematic, global screens for function associated with IDRs that are mapped to protein coordinates: mitochondrial N-terminal targeting signals identified in a proteome-wide screen (Vögtle et al., 2009) and high confidence Cdc28 phosphorylation sites (which point to kinase substrates that are key regulatory sites) curated from *in vitro* and *in vivo* studies (Lai et al., 2012). Importantly, these datasets are unseen by the model during training and testing, and because they are mapped to protein coordinates, they can be used to create IDR-level functional annotations (see Materials and methods). To further ensure that there is no “leakage” between training and test data, we fit the model parameters (*b*, described above) to protein-level annotations from 80% of the proteome, and evaluate the performance on the unseen IDR-level data for the 20% of proteins that were not part of the training dataset. We find that the held-out predictions from FAIDR do as well or better than state-of-the-art predictors of mitochondrial targeting signals and Cdc28 substrates that were specifically designed to predict these functions (Mitofates and Condens, respectively) (Fukasawa et al., 2015; Lai et al., 2012) (Figure 1). This suggests that our general approach is capturing comparable functional information to previous function-specific predictors.

**Figure 1.**
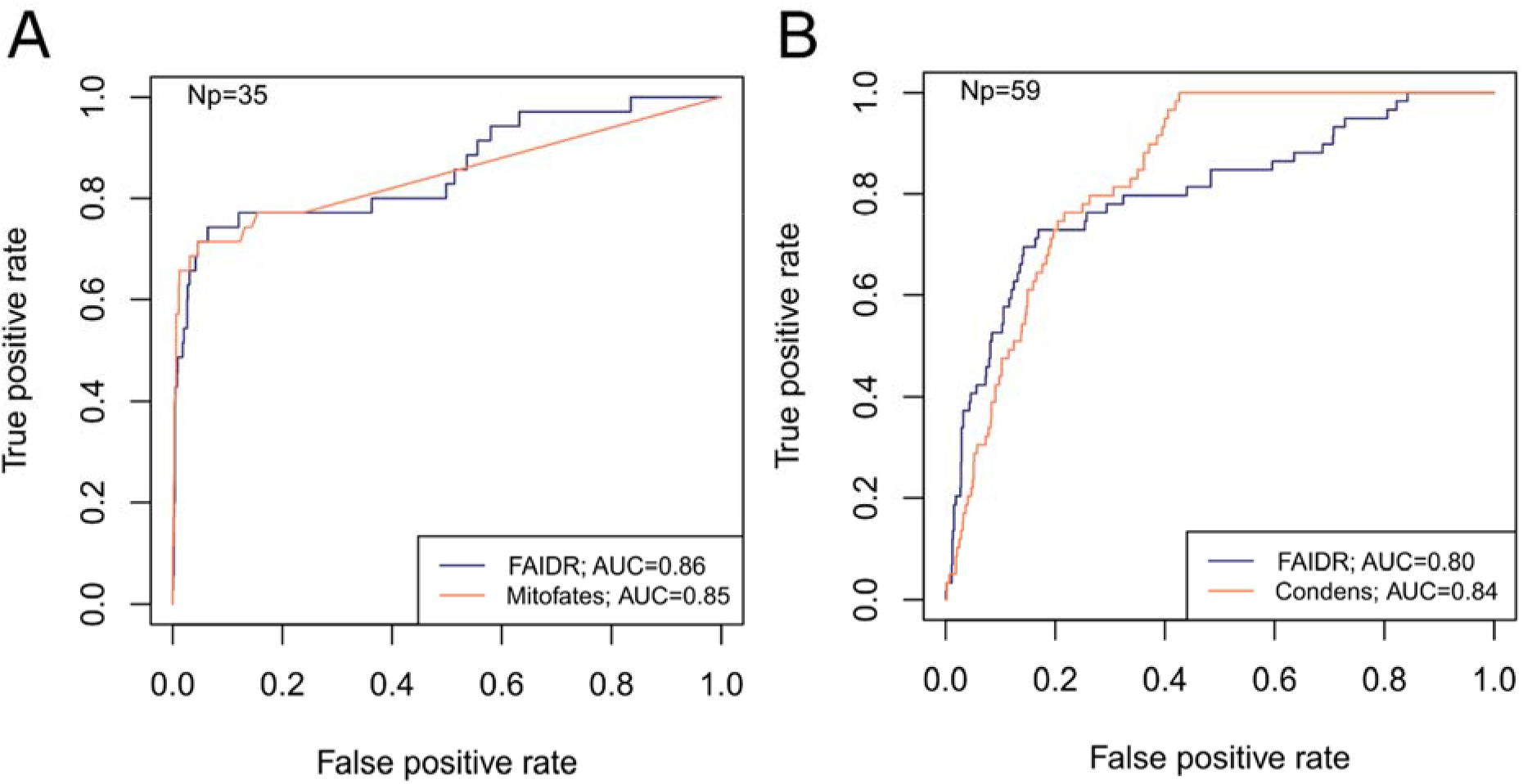
FAIDR trained on evolutionary signatures of IDRs can predict unseen data comparably to state-of-the-art, specific predictors. Receiver Operating Curves (ROC) on a held-out 20% sample are shown for FAIDR trained on evolutionary signatures (blue) versus Mitofates (A) and Condens (B). Area under curve (AUC) is indicated in legend on bottom right for each method. The number of positive IDRs (Np) in the held out 20% is indicated in the top left of each plot.

For functions and phenotypes with no IDR-level data (the vast majority), we tested whether we could identify molecular features predictive of IDR function by evaluating the AUC (Area Under the Curve) in cross-validation on held out proteins (see Methods). We found a diverse set of protein annotations and phenotypes that could be predicted with reasonable power (overall average 5-fold cross validation AUC=0.81, Figure S1, grey bars), ranging from well-characterized IDR functions such as signal transduction and mitochondrial targeting, to underappreciated IDR functions such as ribosome biogenesis and DNA damage (full list of functions in Table S1). Although some of these annotations and phenotypes are correlated, we believe this analysis strongly supports the idea that there is rich functional information in IDR sequences (Zarin et al., 2019), and is evidence that the model is not overfitting.

Because evolutionary signatures can be technically difficult and time consuming to compute (Zarin et al., 2019), we also tested whether FAIDR could predict function from single-species profiles of molecular features. Indeed, we found that the 82 sequence features (Zarin et al., 2019) computed from *S.cerevisiae* IDR sequences alone (scaled to unit variance and 0 mean) could also predict protein function (overall average 5-fold cross validation AUC = 0.77, Figure S1, unfilled bars). Although the overall power was significantly less than the evolutionary signatures (paired t-test over 23 classes p<0.001), for some IDR functions the predictive power was similar, which suggests that IDR function can be predicted from single protein sequences. Identifying informative molecular feature descriptions for IDRs that are easier to compute than evolutionary signatures is a promising area for further research (see Discussion).

### Exploring molecular features of IDRs associated with diverse functions and phenotypes

We next sought to determine which molecular features were associated with IDR function. To do so, we obtained the features that were assigned non-zero coefficients using FAIDR, and fit a standard logistic regression to that subset of features. We extracted the associated t-statistics and visualized them as a clustered heatmap (Figure 2a). Encouragingly, we found that related functions group together in this analysis (Figure 2a, indicated by symbols above function names). For example, consistent with their putative functions (Darling et al., 2017), PMLOs (proteinaceous membrane-less organelles) (Figure 2a, unfilled square) clustered with several phenotypes and functions related to RNA (Figure 2a, filled square).

**Figure 2.**
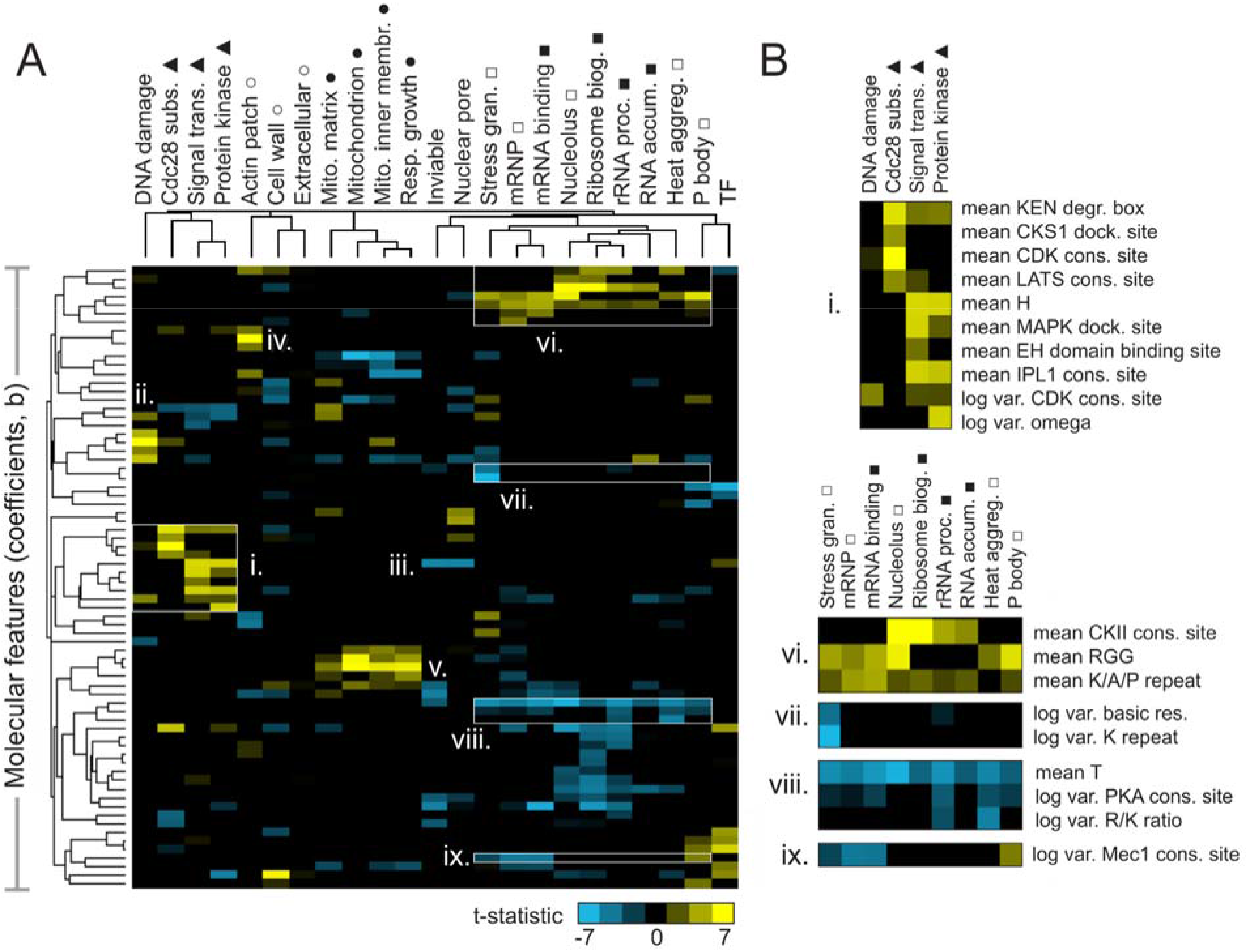
Molecular features of IDRs are associated with specific functions. A) Hierarchical clustering of t-statistics obtained from regression of 23 functions and phenotypes on evolutionary signatures (means and variances of 82 molecular features). Symbols beside function/phenotype names indicate related functions/phenotypes. B) Examples of positive and negatively predictive molecular features for signaling (i) and PMLOs (vi-ix).

We note that the features that underlie these t-statistics represent the extent to which the mean or (log) variance of a sequence-distributed molecular feature (such as a physicochemical property or amino acid repeats) differs from our null expectation over evolutionary time (see Materials and Methods for details). It is relatively straightforward to interpret a positively predictive mean of a property such as hydrophobicity to indicate that elevated hydrophobicity is associated with a certain function. It is more difficult to interpret how the variance of a sequence-distributed property over evolution can be interpreted with respect to its function. For both mean and variance of sequence-distributed properties, we interpret deviations from our null expectation to point to the properties that are (positively or negatively) associated with function (see Discussion).

We found that several molecular features known to be associated with specific biological functions were recovered in this analysis. For example, signaling-associated functions such as Cdc28 kinase substrates, signal transduction, and protein kinase activity share an increase in KEN degradation boxes (Figure 2b, i), while Cdc28 substrates (as expected) are distinguished by increased CKS1 docking sites and CDK consensus sites (Figure 2b, i). The DNA damage response seems to share some features with the more general functions discussed above, but is also (as expected) associated with the presence of specific motifs: PCNA/Pip boxes, Mec1/Tel1 consensus sites (Lai et al., 2012), and NLSs (nuclear localization signals) (Figure 2a, ii). Signal transduction and protein kinase activity share increased histidine residues (Figure 2b, i), which we speculate is related to histidine kinase signaling. Interestingly, the opposite (a decrease in histidine residues) is predictive of essentiality (“inviable” phenotype) and nuclear pore localization (Figure 2a, iii). Other features that are consistent with known disordered region function include the presence of PRK1 consensus sites as a predictor of actin cortical patch function (Figure 2a, iv) (Sekiya-Kawasaki et al., 2003), and increased isoelectric point as a predictor of mitochondrial functions (Figure 2a, v) (reviewed in (Jaussi, 1995)).

We next focused on features predictive of PMLOs, as it is currently unclear whether the association of IDRs with these organelles is determined by amino acid sequence. As expected, we found the presence of RGG (Chong et al., 2018) and K/A/P repeats (Van Der Lee et al., 2014) among the strongest predictors for all these compartments (Figure 2b,vi). We also found a decrease in threonine residues associated with these, as well as all the RNA-related functions (Figure 2b, viii), which to our knowledge has not been previously reported. We were particularly interested in features that could discriminate between PMLOs. We found that stress granules are distinct from all other bodies, showing a decrease in the variance of basic residues and lysine repeats (Figure 2b, vii). Nucleoli and associated ribosomal functions are distinct in that they show increased CKII consensus sites (Figure 2b, vi). We also found that processing body (P body) proteins show an increased variance in Mec1 consensus sites, while mRNPs and mRNA binding proteins show a decrease in this feature (Figure 2b, ix).

### Molecular features in the Cox15 IDR predictably affect mitochondrial targeting of Cox15

In order to directly test whether the features in our evolutionary signatures that are predictive of function are required for function (and not simply correlated with function), we sought to manipulate these features in an example IDR. For this, we chose the N-terminal IDR of the Cox15 protein, which is a known mitochondrial targeting signal (Vögtle et al., 2009), and which we have previously used to explore phenotypic consequences of different IDR genotypes (Zarin et al., 2019). Using FAIDR, we first identified the top predictive features for this IDR (Cox15 IDR 1, which spans amino acids 1 through 45 in Cox15) that indicate association with mitochondrial localization (“mitochondrion” GO annotation), and plotted the feature vector (evolutionary signature) values corresponding to these top features in Figure 3A. The top predictive feature (a positive predictor) is mean isoelectric point (pI), which also has a relatively high Z-score in the Cox15 IDR 1 feature vector. This indicates that increased isoelectric point is not just a generally positive predictor for mitochondrial localization, but that elevated isoelectric point is present in the Cox15 IDR specifically. Another top positive predictor that is present in this IDR is increased hydrophobicity (Figure 3A).

**Figure 3.**
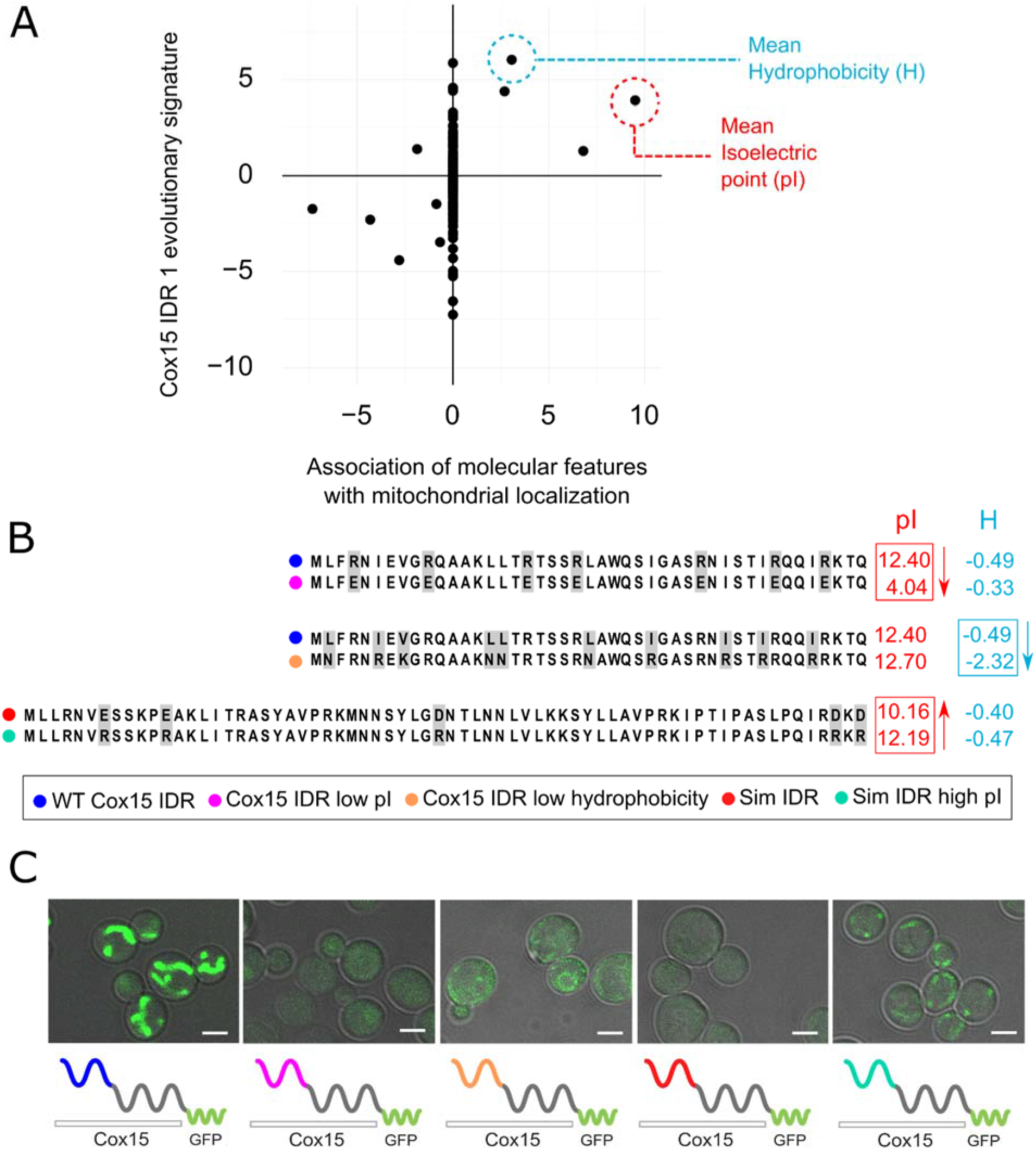
Molecular features predicted to be associated with “mitochondrion” GO annotation affect mitochondrial localization phenotype. A) Feature vector (evolutionary signature) of the Cox15 N-terminal IDR (Cox15 IDR 1) (y-axis) is plotted against predictive features for mitochondrial function as determined by FAIDR (x-axis). Two of the top features associated with mitochondrial localization are isoelectric point (pI), circled in red, and hydrophobicity (H), circled in blue B) Amino acid sequences for each Cox15 IDR variant. Wildtype and simulated Cox15 IDR sequences are compared to the same sequence with mutations altering isoelectric point (pI) and hydrophobicity. Sequences with variable residues (grey) are visualized with Jalview (Waterhouse et al., 2009) C) Micrographs showing the mitochondrial localization phenotype for different budding yeast strains that differ in their Cox15 N-terminal IDRs. Green shows GFP-tagged Cox15 localization. Left to right: wildtype IDR, Cox15 IDR with low pI, Cox15 IDR with low hydrophobicity, simulated IDR, simulated IDR with high pI. Scale bar represents 1 μm.

In order to manipulate these sequence-distributed features (isoelectric point and hydrophobicity), we constructed strains with different Cox15 IDR 1 genotypes and molecular features (Figure 3B). We first created a mutant IDR with lower pI by mutating each arginine residue (R) in the wildtype IDR sequence to a glutamic acid (E), which lowered the pI of the wildtype IDR from 12.40 to 4.04 while only minimally changing the hydrophobicity (Figure 3B, top). As expected, the wildtype GFP-tagged Cox15 strain has a clear mitochondrial localization (Figure 3C, left). However, the “Cox15 IDR low pI” strain no longer localizes to the mitochondria.

This provides evidence that the elevated pI is necessary for mitochondrial targeting function. Similarly, we created a mutant IDR with lower hydrophobicity by mutating each leucine residue (L) to asparagine (N), each valine (V) to lysine (K), and each isoleucine residue (I) to arginine (R) (Figure 3B, middle). This changed the three most hydrophobic residues into the three least hydrophobic according to the Kyte-Doolittle scale (Kyte and Doolittle, 1982), while only minimally affecting the isoelectric point. Interestingly, the “Cox15 IDR low hydrophobicity” strain also fails to localize to the mitochondria (Figure 3C), strongly supportive of elevated hydrophobicity also being necessary for mitochondrial targeting function.

To test if elevated pI is sufficient to restore a targeting signal, we first constructed an N-terminal Cox15 IDR in which we “knocked out” sequence-distributed features that we predict to be important for specific IDR function. To make this synthetic Cox15 IDR, we simulated the evolution of the Cox15 N-terminal IDR under a model that randomly evolves intrinsically disordered regions while preserving position-specific variation in evolutionary rates, but includes no specific constraints on the sequence-distributed molecular features used to train FAIDR (as in (Iserman et al., 2020)) (Figure 3B, bottom). We note that this simulated IDR is not a completely random or generic IDR; for example, the simulated IDR shares 4/5 of the N-terminal amino acids with the wildtype Cox15 IDR from which it “evolved” *in silico* (Figure 3B). Nevertheless, consistent with the functional importance of the sequence-distributed features, the strain in which the N-terminal IDR of GFP-tagged Cox15 is replaced with this simulated IDR fails to show a mitochondrial localization pattern (Figure 3C), indicating that the first few conserved amino acids in the IDR are not sufficient for mitochondrial localization. We next made mutations in the simulated Cox15 IDR to increase the pI back to the range of the wildtype Cox15 N-terminal targeting signal. We mutated each glutamic acid (E) and aspartic acid (D) residue to arginine (R), bringing the pI from 10.2 in the simulated IDR to 12.2 in the simulated IDR with high pI. Remarkably, this mitochondrial localization phenotype seems to be partially rescued when the pI of the simulated IDR is increased (“sim IDR high pI”, Figure 3C). This indicates that the elevated pI in the context of a synthetic, non-functional Cox15 N-terminal disordered region is sufficient to restore weak mitochondrial targeting function.

Taken together, these results demonstrate that functional associations with evolutionary signatures revealed by the FAIDR statistical model can be used to design mutations in IDRs that alter phenotype in predictable ways, and suggest that a small number of molecular features may be necessary and sufficient for partially restoring some IDR functions.

### Inferring function for specific IDRs in uncharacterized proteins

One of the important use-cases of our approach (the FAIDR model trained on evolutionary signatures of predicted IDRs) is to predict functions for specific protein regions that are otherwise challenging, if not impossible, to annotate using current methods. To demonstrate the use of FAIDR for annotating protein regions, we chose to focus on open reading frames (ORFs) annotated as “uncharacterized” in the yeast proteome by SGD (Cherry et al., 2012). There are currently 739 uncharacterized ORFs in the yeast proteome, 121 of which have one or more IDRs for which we are able to make predictions. We extracted the probability that each IDR corresponds to each of the functions that we considered (equation 1), and used this to make functional predictions for IDRs in uncharacterized ORFs.

One interesting example is the N-terminal IDR in putative mitochondrial protein SHH3, which we predict to be associated with the mitochondrion, as well as the mitochondrial inner membrane. Indeed, SHH3 assumes what looks to be a mitochondrial localization in recent localization collections such as the LoQAtE (Localization and Quantitation Atlas of the yeast proteomE) database (Breker et al., 2014, 2013) and Yeast RGB (Dubreuil et al., 2019). Another interesting case is an IDR in MFG1, which FAIDR predicts to be associated with sequence-specific DNA binding (Figure 4A), indicating that MFG1 could be a transcription factor. Indeed, all other IDRs with a higher probability for association with transcription factor activity according to FAIDR are IDRs from confirmed transcription factors (Sok2, Sfp1, Mot3) (Figure 4A). Despite being an uncharacterized ORF, Mfg1 regulates filamentous growth by interacting with the FLO11 promoter and regulating FLO11 expression (Ryan et al., 2012), and has a nuclear localization (Huh et al., 2003), further supporting the idea that it is indeed a transcription factor. Using FAIDR, we can specifically identify the Mfg1 protein region that is associated with transcription factor activity (Figure 4B). Furthermore, by looking at the top predictive features that FAIDR points to (Figure 4C, left) and plotting the feature vector (evolutionary signature) for these top features, we can begin to understand the molecular features that are contributing to this function for the Mfg1 IDR 1 specifically. For example, increased variance of glutamine residues in Mfg1 is a positive predictor of its association with transcription factor activity (Figure 4C, orange box), and this sequence feature can be seen in a multiple sequence alignment of the Mfg1 IDR and its orthologs (Figure 4C, right). Thus, we can formulate the hypothesis that this specific sequence feature is important for function, and test this by modulating the number of glutamine repeats in the lab (see Discussion).

**Figure 4.**
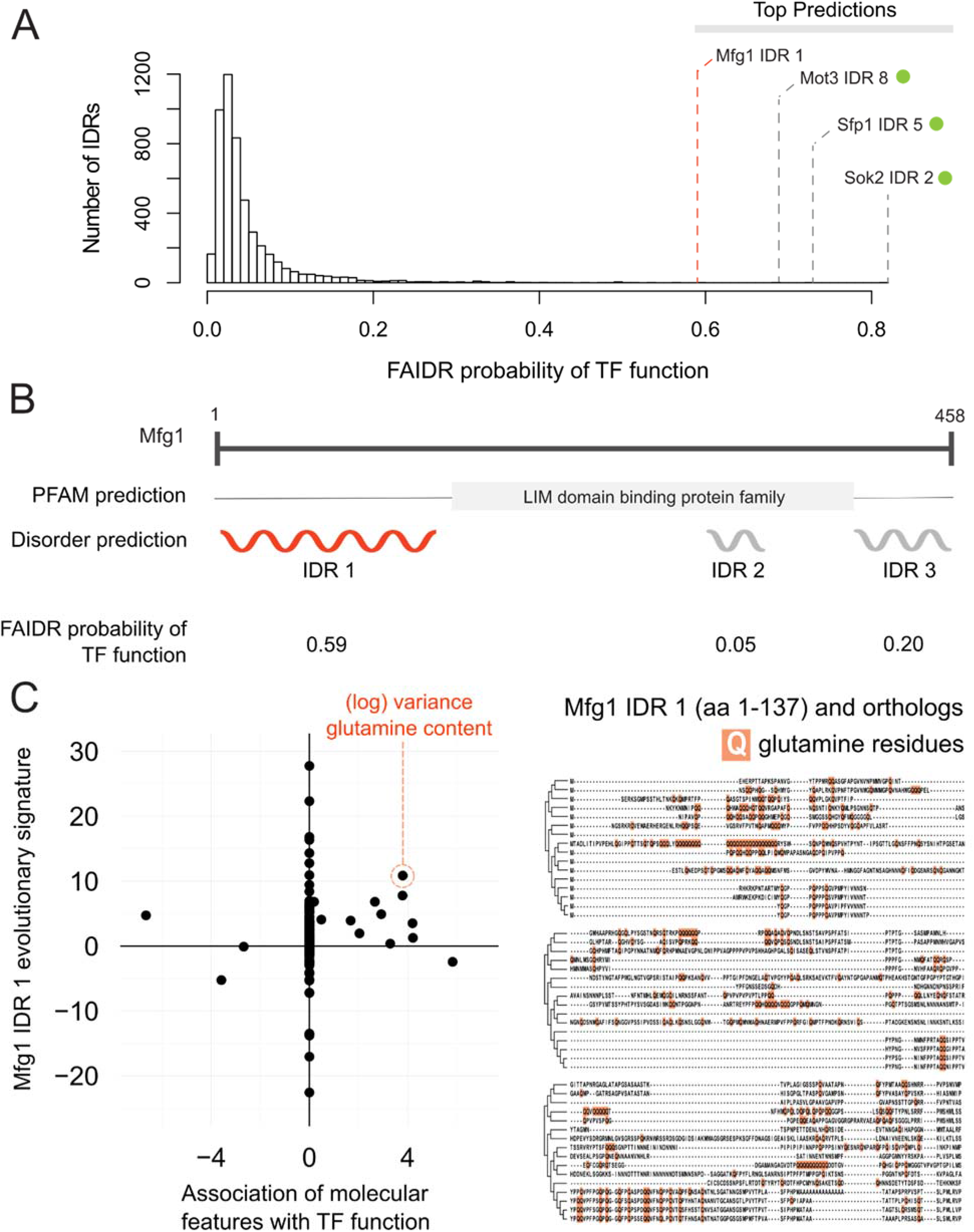
Identification of a specific IDR sequence associated with transcription factor activity in an uncharacterized protein. A) Histogram of probabilities for association of IDRs with sequence-specific DNA binding (transcription factor, or “TF”) function. Top predictions are indicated. From top predictions, IDRs from known transcription factors are indicated with green dot. Mfg1 is an uncharacterized protein. B) Mfg1 protein coordinates, with all known domain annotations (via PFAM), disorder prediction (via DISOPRED3), and FAIDR probability for association with sequence-specific DNA binding indicated. C) Feature vector (evolutionary signature) for Mfg1 IDR 1 (y-axis) is plotted against predictive features for transcription factor/sequence-specific DNA binding function as determined by FAIDR (x-axis). Log variance in glutamine (Q) residues is highlighted as one of the top features (orange circle) associated with TF function. Glutamine residues (Q) are indicated (in orange) in the IDR of Mfg1 and orthologs obtained from the YGOB (Byrne and Wolfe, 2005). Multiple sequence alignment of orthologs is visualized using Jalview (Waterhouse et al., 2009). Species in alignment (from bottom to top) are: *Saccharomyces cerevisiae, Saccharomyces mikatae, Saccharomyces kudriavzevii, Saccharomyces uvarum, Candida glabrata, Kazachstania africana, Kazachstania naganishii, Naumovozyma castellii, Naumovozyma dairenensis, Zygosaccharomyces rouxii, Torulaspora delbrueckii, Eremothecium (Ashbya) gossypii, Eremothecium (Ashbya) cymbalariae, Lachancea kluyveri, Lachancea thermotolerans, Lachancea waltii.*

## Discussion

We have demonstrated a systematic approach to associate molecular features of IDR sequences with diverse biological functions. Our method stands in contrast to those established for specific subtypes of IDRs, as well as existing methods that use amino acid sequences to predict whole protein function. Our goal is to generate hypotheses both about the function of individual IDRs and which molecular features in those IDRs are important for those functions, much like Pfam annotations provide both predictions of molecular functions of domains, as well as hypotheses about specific residues within those domains that are important for that function (El-Gebali et al., 2019).

In the case of Mfg1, which is a protein of unknown function, the N-terminal IDR is among the IDRs most strongly associated with transcription factor function in the yeast proteome. Because Mfg1 contains three predicted IDRs (and in general proteins contain multiple IDRs), this illustrates the advantage of our approach that infers which (if any) of the IDRs are associated with a particular function. Although in some well-studied cases, it is clear that the C-terminal and N-terminal IDRs have different functions (e.g., p53, (Krois et al., 2018; Laptenko et al., 2016)), in general we do not know if functional information within IDRs is modular (confined to one IDR) or distributed (spread over all the IDRs in a protein). In our exploration of yeast IDRs strongly associated with functions, we identified interesting cases where multiple IDRs within a protein were strongly associated with different functions (e.g., Supplementary figure S3) possibly supporting the idea that IDRs can be thought of as functional modules. However, we note that the identification of IDR boundaries in protein sequences (via bioinformatics or experiments) is an error-prone process (Nielsen and Mulder, 2019) and thus in many cases we are not confident how many IDRs a protein ‘truly’ contains. A key technical challenge hindering inference of the IDR sequence features that are associated with function is that most proteome-scale functional information is at the level of entire proteins and proteins often contain multiple predicted IDRs, the exact boundaries of which are often uncertain. Further improvement of the approaches described here could allow simultaneous inference of IDR function and boundaries, perhaps providing more insight.

The analysis of Mfg1 also reveals that homologues of the N-terminal IDR show elevated variance in glutamine repeats, a property of IDRs that we find associated with transcription factor function, consistent with previous reports (Freiman and Tjian, 2002; Gemayel et al., 2015; Klemsz and Maki, 1996). Thus, in addition to providing a hypothesis about which IDR is important for Mfg1 function, we predict that modulation of the glutamine residues in the N-terminal IDR of Mfg1 will impact its function in transcription. We note that in the case of Mfg1, the molecular feature is an elevation of evolutionary variance, and this feature cannot be modulated experimentally because we only affect one single value. Instead, we interpret the associations with our evolutionary signatures as pointing towards testable hypotheses. While FAIDR analysis using descriptions of molecular features based on single sequences would be easier to interpret, we found that our features did not perform as well when computed on *S. cerevisiae* alone (Figure S1). This supports the use of evolutionary features in this case, but as we discuss further below, suggests that development of improved molecular features that can be manipulated in single sequences is an area for future research.

In addition to other previously reported molecular features (the expected short linear motifs and repeats associated with signaling, posttranslational regulation and RNA binding) we found an unexpected depletion of threonine residues in IDRs in most classes of RNA-binding proteins and an unexpected decreased variance of lysine residues specific to stress-granule proteins. Future experiments will test the importance of these features. Nevertheless, our interpretable molecular features and statistical framework both confirms previous associations with function and reveals new properties of IDRs, while still providing predictive power comparable to current function-specific bioinformatics approaches.

At least in the case of the Cox15 N-terminal mitochondrial targeting signal, we could show that two molecular features, mean isoelectric point and mean hydrophobicity, the increase of which we found to be positively associated with function, appear to be necessary for function of that IDR (Figure 3). To facilitate wider use of this approach, we have developed a website that provides search, visualization and comparison of evolutionary signatures.

Remarkably, we also found that artificially increasing the isoelectric point of a simulated Cox15 N-terminal IDR (i.e., an IDR with no enrichment of sequence-distributed features) enabled us to create a weak synthetic targeting signal. To our knowledge this represents the first synthetic IDR designed to perform a specific biological function, and highlights the benefit of using a simple, interpretable statistical model: we could hand-design mutations in a synthetic IDR to perform a desired function through 8 amino acid changes. In general, designing synthetic IDRs represents a new approach for testing the importance of molecular features.

In all, we found that diverse functions can be associated with the IDR sequence features included in our evolutionary signatures (Zarin et al., 2019). However, we note that these features of disordered regions were obtained from a review of the literature, and new features are continuously being discovered (e.g., patterning of aromatic residues (Martin et al., 2020)). It is likely that including these features will further improve predictive power of the approach described here. Nevertheless, these features are necessarily biased by current research interests, and because they rely on comparisons to simulations of molecular evolution, they are cumbersome to compute. In future, we believe application of advanced computational methods will enable more systematic exploration, ideally automatically learning molecular features that are most important for IDR function.

## Materials and methods

### The FAIDR statistical model

FAIDR was implemented in R with the glmnet package (Friedman et al., 2010). Code and results are available at http://www.github.com/taraneh-z/FAIDR. A schematic of the model is provided in Supplementary materials (Fig. S2).

To maximize the likelihood of our model (eq. 2) we use the E-M algorithm, or iteratively maximize the expected complete log likelihood (Moses, 2017). To derive the E-M algorithm for this model, we first write the complete likelihood (*CL*, the log likelihood assuming the hidden variables are observed):

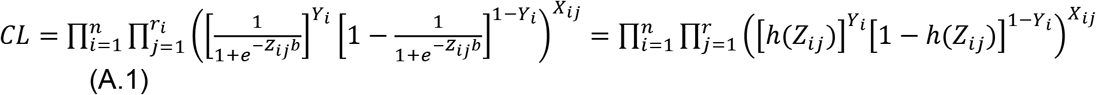

where 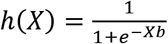

Taking logs and expectations yields:

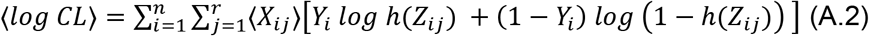

Where the angle brackets denote expectations. In the E-step of the E-M algorithm, these expectations are calculated using Bayes’ theorem as follows:

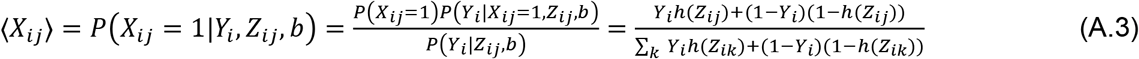

Note that in (eq. A.3) we have assumed no prior knowledge about which IDR in a protein is most likely to be responsible for the function (uninformative priors), *P*(*X_ij_* = 1) is constant over j.

In the M-step, we maximize the expected complete log likelihood. Inspection of equation A.2 reveals exactly one term for each IDR, weighted by 〈*X_ij_*〉, the expected value of the hidden variable. Thus, our model corresponds to an iteratively reweighted logistic regression problem, where the weights are the current guess of the hidden variable, 〈*X_ij_*〉, representing which IDR is responsible for the function. We can therefore maximize the L1 penalized version of this expected complete log likelihood by adding an extra factor to the weights of the IRLS algorithm (Hastie et al., 2015b). The modified weights are:

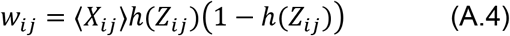

In each M-step in our implementation, we do 5 iterations of this IRLS (on standard LASSO regression) to obtain new parameter values. We used and a fixed L1 penalty of 0.2 throughout, as we found this to provide the strong regularization needed for our small numbers of positive examples. To improve the numerical stability of IRLS (needed when we had small numbers of positive examples relative to the number of features) we also used a small L2 penalty (alpha =0.99 in glmnet) and we monitored the weights (A.4) and set them to an arbitrarily small value if they reached 0. After 5 iterations of IRLS, we recalculate the expected values of the hidden variables 〈*X_ij_*〉. We initialize 〈*X_ij_*〉 by setting them equal to 1/*r_i_*, so that they are downweighted equally.

### Obtaining evolutionary signatures for IDRs in the yeast proteome

In order to calculate evolutionary signatures from predicted IDRs (disordered regions predicted from the Saccharomyces cerevisiae proteome using DISOPRED3 (Jones and Cozzetto, 2015) that are 30 amino acids or longer), we used a similar method to previously published work (Zarin et al., 2019) with some modifications.

Briefly, for each set of orthologous IDRs (obtained from the Yeast Gene Order Browser (Byrne and Wolfe, 2005) and filtered to include those proteins for which there are 10 or more orthologs), we simulated 1000 sets of orthologous IDRs (using previously described methods (Nguyen Ba et al., 2014, 2012), and quantified the difference in the evolution of molecular features in the real set of orthologous IDRs to that of the simulated orthologous IDRs using a standard Z-score (as described in (Zarin et al., 2019)). The evolutionary signature of each IDR is thus comprised of a vector of 164 Z-scores (mean and log variance over evolution for 82 molecular features (listed in (Zarin et al., 2019)).

In previous work (Zarin et al., 2019), we calculated these evolutionary signatures for highly diverged segments of IDRs by constraining the evolution of short conserved segments in our simulations of IDR evolution. For this study, we wanted to include all of the potentially functional sequence information in our evolutionary signatures (i.e., highly diverged segments as well as short conserved segments), and therefore did not constrain the evolution of short conserved segments in the evolutionary simulations. The evolutionary simulation software (Nguyen Ba et al., 2014, 2012) takes as input maximum-likelihood estimates for the column (site-specific) rate of evolution, the local rate of evolution, and whether or not a site is predicted to be part of a conserved Short Linear Motif (SLiM) (Nguyen Ba et al., 2012). In order to ignore the constraint on these short conserved segments, we randomly scrambled the column (site-specific) rate of evolution for each IDR, and set each found motif to 0. This results in evolutionary simulations of IDRs in which positionally-conserved sites evolve with the same average constraint as other sites in the IDR.

Evolutionary signatures are available for download and visualization at www.moseslab.csb.utoronto.ca/idr/.

### Compiling functions and phenotypes for prediction

We compiled a series of functional annotations, phenotypes, and datasets to predict with our model. These fell into the following categories:

1. Gene Ontology annotations that we previously found to be strongly enriched in our unsupervised clustering analysis (Zarin et al., 2019), acquired from SGD (Cherry et al., 2012).
2. Datasets that screened the S.cerevisiae proteome for membraneless organelle/reversible protein assembly formation under stress (Mitchell et al., 2012; Narayanaswamy et al., 2009; Wallace et al., 2015).
3. A dataset of gold-standard Cdc28 substrate predictions (Sharifpoor et al., 2011).
4. A set of publicly available S.cerevisiae phenotype annotations from SGD (Cherry et al., 2012).

In total we found 23 functions and phenotypes for which we could get accurate predictions (Figure S1). Overall, we tested the model on 120 functions and phenotypes.

### Evaluating FAIDR at the IDR level

We obtained two datasets for which we could define “positive” and “negative” IDRs from proteomics data.

First, we obtained a dataset of mitochondrial targeting signals from a global analysis that identified and validated mitochondrial targeting signals in highly purified mitochondria using combined fractional diagonal chromatography (COFRADIC) and other methods (Vögtle et al., 2009).

Second, we obtained Cdc28 phosphorylation sites discovered in high-throughput kinase-specific MS experiments (Holt et al., 2009; Lai et al., 2012) and mapped them to intrinsically disordered regions. If at least one verified phosphorylation site was identified in a region, we considered this intrinsically disordered region to be a target of Cdc28.

To make predictions of functions for unseen IDRs, we used the parameters (*b*) estimated for each function and computed equation (1) above. We compared these probability values to the known IDR function as the probability cutoff is varied. We used ROCR (Sing et al., 2009) to compute ROC curves and AUCs.

Even though these datasets were not used directly for the protein-level annotations, they may have been used as part of the evidence that led to the annotation of the protein. Therefore, we sought to further remove the potential for circularity by removing 20% of proteins from the training dataset and only evaluating the predictive power on the IDRs from those 20%. We report the number of positive IDRs in this held out data set in Figure 1.

### Evaluating FAIDR at the protein level

Although predicting protein function was not a primary goal of FAIDR, we evaluated the AUCs on held out protein annotations to confirm that FAIDR could learn features that are predictive of a diverse set of IDR functions. To make predictions of functions for held out proteins (Y), we sum the weighted contributions of all the IDRs:

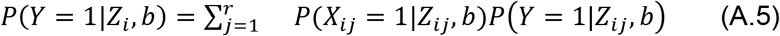

where *P*(*Y* = 1|*Z_ij_, b*) is given in equation (1) above. To compute *P*(*X_ij_* = 1|*Z_ij_, b*) for an unseen sequence, we cannot use (A.3) because it treats *Y* as an observed variable. Instead we used: 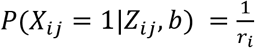, which amounts to a simple average over the IDRs in each protein.

For each functional category/annotation we used 5-fold cross-validation, randomly dividing the data into 80% of proteins for training and 20% of proteins for testing. For each fold of cross-validation, we compute the AUC using ROCR (Sing et al., 2009) on the random 20% held out data. We report the average AUC over the 5 held out datasets.

### Clustering of t-statistics

We fit a standard logistic regression to molecular features that obtained non-zero coefficient values from FAIDR. We extracted the corresponding t-statistic for each molecular feature, and clustered these t-statistics and the 23 functions that we predicted using hierarchical clustering for Figure 2. Prior to clustering, we assigned those molecular features with a coefficient of 0 a corresponding t-statistic of 0, and filtered the 164 molecular features to only include those molecular feature vectors that had at least one t-statistic value of 3 or greater for a predicted function. This filtering resulted in a total of 72 molecular features. We used the Cluster 3.0 program (de Hoon et al., 2004) to perform hierarchical clustering (clustering both “genes” [molecular feature t-statistics] and “arrays” [functions] using uncentred correlation distance, with calculated weights and using average linkage). We visualized the clustering using Java Treeview (Saldanha, 2004).

### Construction of Cox15-GFP strains with variable molecular features and confocal microscopy

We obtained a BY4741-derived Cox15-GFP tagged budding yeast strain from the Yeast GFP collection (Huh et al., 2003), which serves as our wildtype IDR strain. We mutated the N-terminal IDR in this strain as previously described (Zarin et al., 2019) using the Delitto Perfetto method (Storici et al., 2001), which performs markerless site-directed mutagenesis in the genome. In order to construct the IDR mutant strains (“Cox15 low pI”, “Cox15 low hydrophobicity”, “Sim IDR”, and “Sim IDR high pI”), we used gene synthesis (via Integrated DNA Technologies [IDT]) whose sequences we confirmed using PCR and Sanger sequencing. The genotypes for the IDR strains are as follows:

WT:

ATGCTTTTCAGAAACATAGAAGTGGGCAGGCAGGCAGCTAAGCTATTAACGAGAACCTCGAGTCGTTTGGCCTGGCAAAGTATTGGGGCCTCAAGGAATATTTCTACCATCAGACAACAAATCAGAAAGACTCAA

Cox15 low pI:

ATGCTTTTCGAAAACATAGAAGTGGGCGAACAGGCAGCTAAGCTATTAACGGAAACCTCGAGTGAATTGGCCTGGCAAAGTATTGGGGCCTCAGAAAATATTTCTACCATCGAACAACAAATCGAAAAGACTCAA

Cox15 low hydrophobicity:

ATGAATTTCAGAAACAGAGAAAAGGGCAGGCAGGCAGCTAAGAATAATACGAGAACCTCGAGTCGTAATGCCTGGCAAAGTAGAGGGGCCTCAAGGAATAGATCTACCAGAAGACAACAAAGAAGAAAGACTCAA

Sim IDR:

ATGCTGCTGAGAAACGTTGAATCCTCCAAACCCGAAGCAAAACTAATTACCAGAGCTTCTTACGCCGTGCCCAGGAAAATGAACAATTCATACTTGGGCGATAATACATTGAATAACCTGGTCTTAAAGAAGAGCTATCTTTTAGCTGTTCCCAGAAAGATTCCCACGATTCCAGCCAGTCTGCCGCAAATTCGTGACAAGGAT

Sim IDR high pI:

ATGCTGCTGAGAAACGTTAGATCCTCCAAACCCAGAGCAAAACTAATTACCAGAGCTTCTTACGCCGTGCCCAGGAAAATGAACAATTCATACTTGGGCAGAAATACATTGAATAACCTGGTCTTAAAGAAGAGCTATCTTTTAGCTGTTCCCAGAAAGATTCCCACGATTCCAGCCAGTCTGCCGCAAATTCGTAGAAAGAGA

Confocal Microscopy (Leica TCS-SP8, 63X oil immersion objective) was used to image cells. Briefly, log phase cultures were spun down at 3000 rpm for 3 minutes to concentrate cells, after which live cells were placed on standard, uncoated glass slides for imaging. Cells containing GFP-tagged proteins were excited with a 488 nm laser. Each strain was imaged on at least three separate days. Images in Figure 3C were taken on the same day with the same microscope settings across all images.

## Acknowledgements

We thank Alex X. Lu for comments on the manuscript and Dr. Alan R. Davidson for discussions about IDR functional predictions. We thank Dr. Helena Friesen and Dr. Brenda Andrews for providing strains from the yeast GFP collection. We thank Canadian Institutes for Health Research (CIHR) for funding to AMM and JDF-K (grant no. PJT-148532), the Canada Research Chairs program and a CIHR Foundation grant (grant no. FDN-148375) to JDF-K, Canada Foundation for Innovation (CFI) for funding to AMM, and the Natural Sciences and Engineering Research Council of Canada (NSERC) for funding to TZ.

## Supplementary materials

**Table S1.**
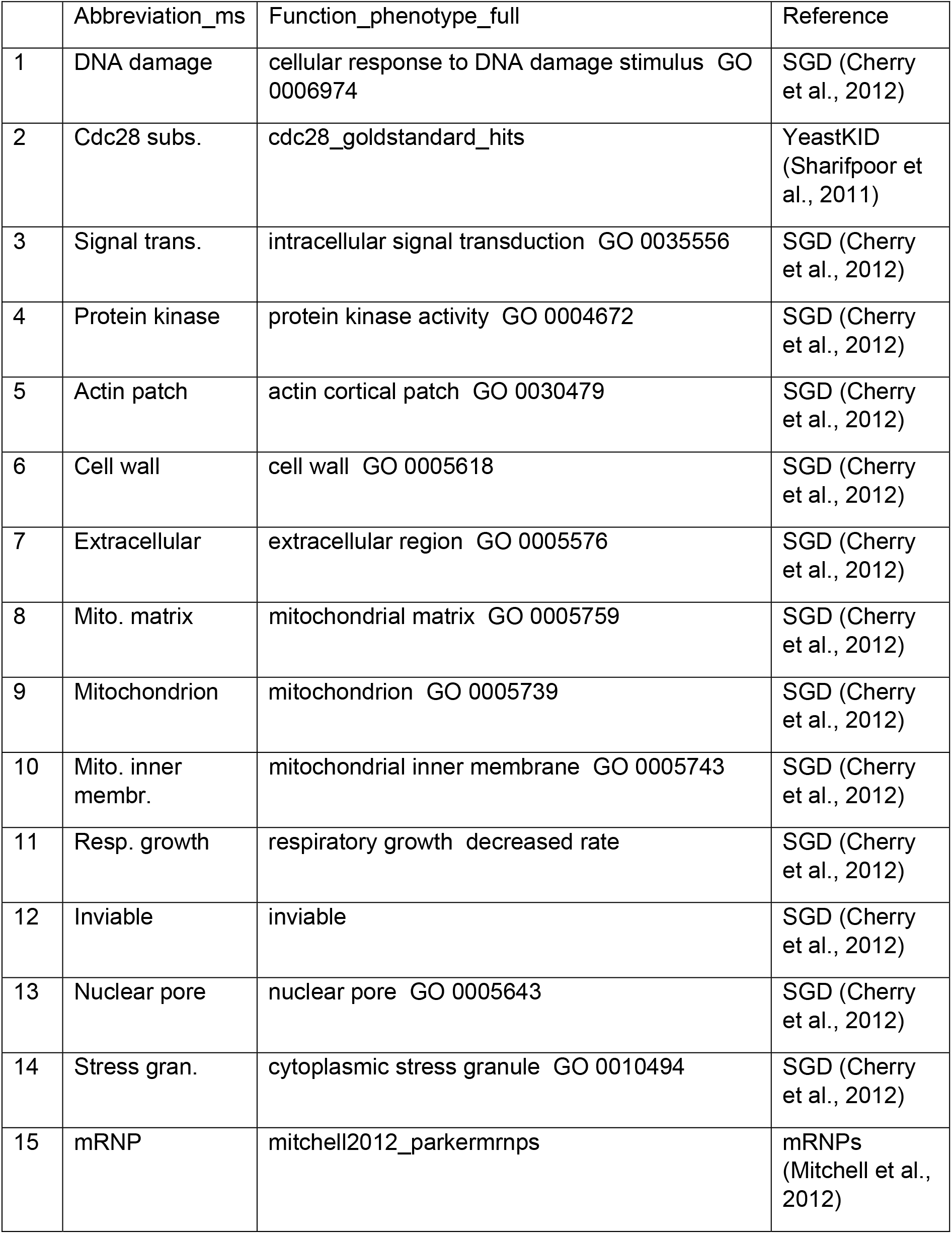
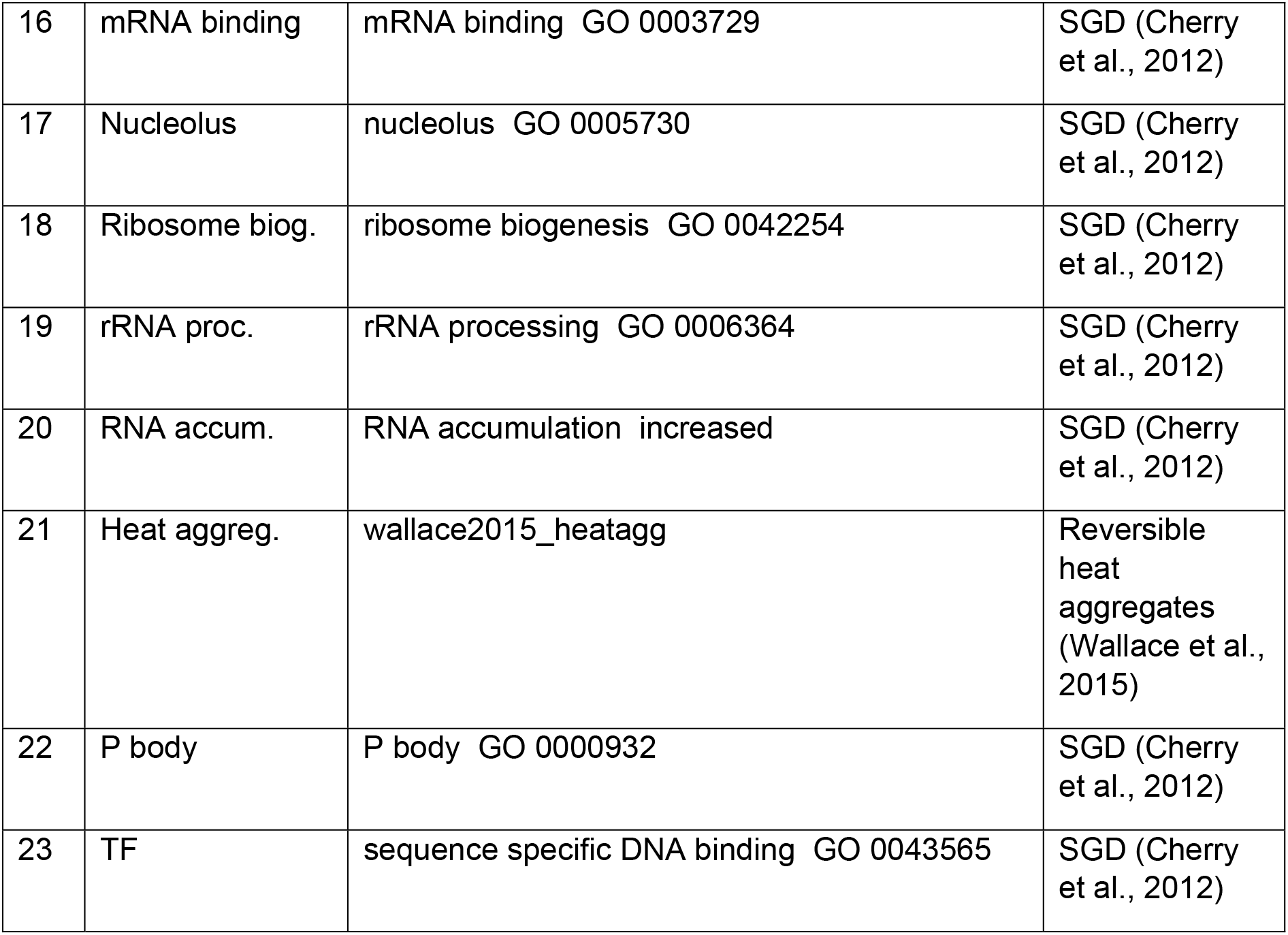
Table of function and phenotype names, abbreviations, and references.

**Figure S1.**
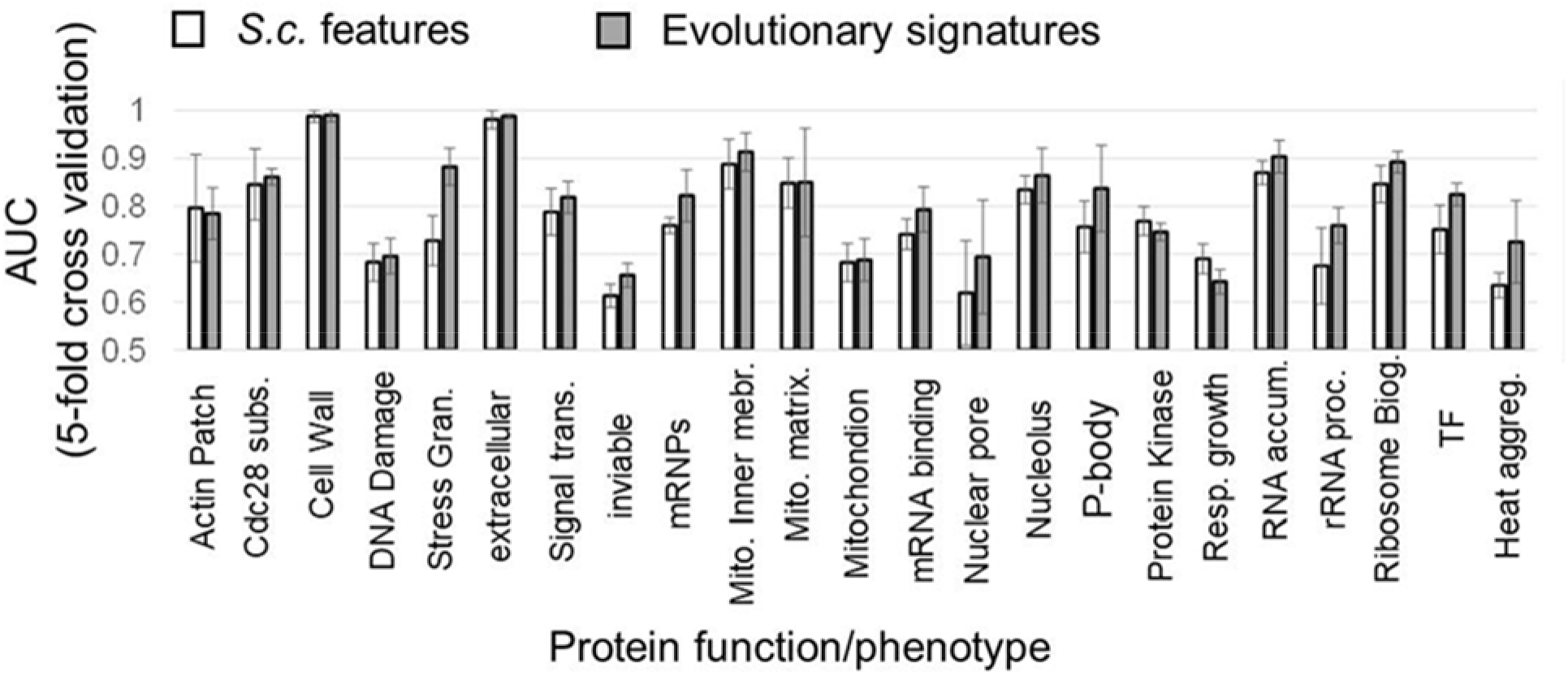
Predicting diverse protein function and phenotype using FAIDR. Average Area under the Receiver Operating Curve (AUC, computed using ROCR (Sing et al., 2009) on the held out 20% in 5-fold cross-validation is shown, computed using either the 82 molecular features on the S. cerevisiae IDR sequence alone (unfilled bars), or from evolutionary signatures (filled bars, see text). Error bars represent the standard deviation. Functional annotations are from GO (obtained from SGD (Cherry et al., 2012)), except gene deletion phenotypes (inviable, respiratory growth, RNA accumulation, from SGD), Cdc28 substrates (yeastKID, (Sharifpoor et al., 2011)), mRNPs (Mitchell et al., 2012)) and reversible heat aggregation (Wallace et al., 2015).

**Fig. S2.**
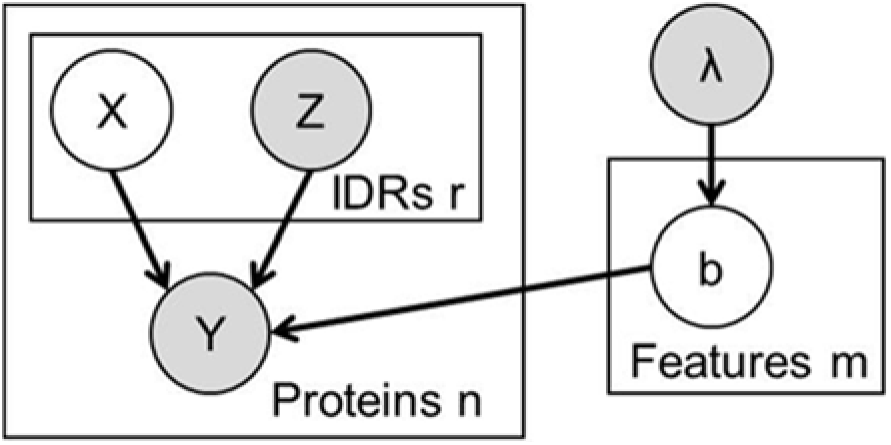
The FAIDR probabilistic model. Unfilled circles represent hidden variables, grey circles represent known variables. Functions or phenotypes are treated as binary variables, where *Y*=1 if the protein is associated with that function or phenotype, and *Y*=0 if the protein is not. For the *i*-th protein, we have a set of features, *Z_i_* for each of *r_i_* disordered regions. We use a hidden variable, *X_i_*, to represent the IDR that is responsible for the function, so that if *X_ij_*=1, the *j*-th IDR in the *i*-th protein is the one responsible for the function. Throughout we used an exponential prior, λ=0.2 on the coefficients, *b*, (equivalent to a constant L1 penalty) that we found to limit overfitting for our functions with small numbers of positives.

**Fig. S3.**
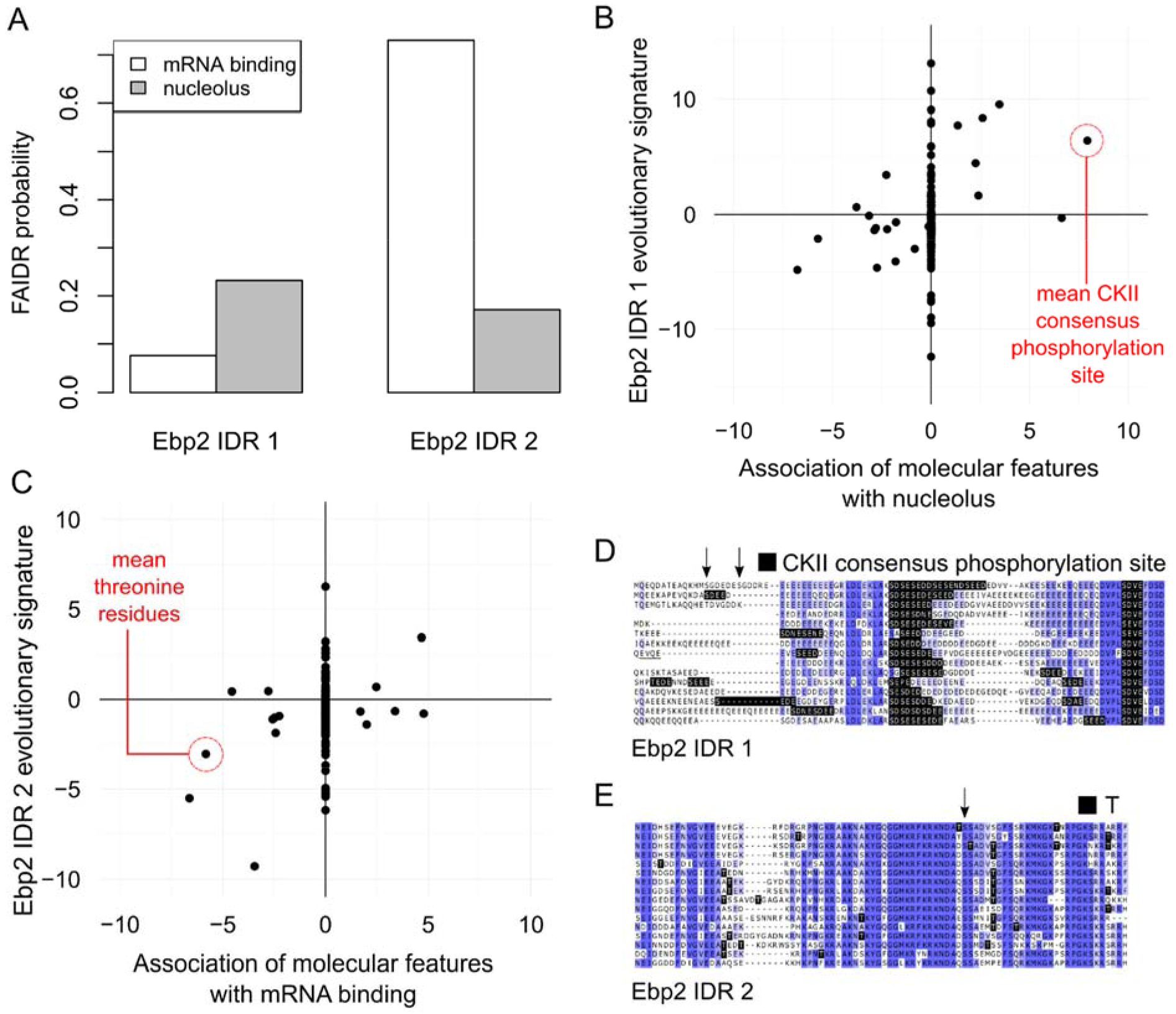
Example of modular IDR function predicted by FAIDR A) FAIDR probabilities for the IDRs in Ebp2. B) and C) show feature vectors (evolutionary signatures) for Ebp2 IDRs (y-axis) plotted against predictive features for nucleolus (B) and mRNA binding (C) (x-axis) as determined by FAIDR. Top predictive features for nucleolus (B) and mRNA binding (C) are highlighted in red. D) and E) show alignments of IDRs from Ebp2, where CKII consensus phosphorylations sites ([ST].[DE][DE]) and (lack of) threonine (T) are indicated by black highlighting, respectively. Arrows above the sequence indicate residues reported to be phosphorylated (from SGD (Cherry et al., 2012)) in S.cerevisiae (top protein sequence in each case). Shade of blue represents percent identity in the multiple sequence alignment (darker blue is higher percent identity). Ebp2 is annotated in GO (The Gene Ontology Consortium, 2019) to be localized to the nucleolus and to bind mRNA. D) and E) show that the first IDR reveals an abundance of CKII consensus phosphorylation sites, while the second IDR reveals reduction in threonine (T). Remarkably, these predictions are consistent with the report that a C-terminal truncation mutant of Ebp2 including the second IDR (Ionescu et al., 2004) leads to defects that are distinct from the nucleolar rRNA related functions. This example illustrates how different IDRs in multi-IDR proteins can be associated with different functions: the IDRs contain different molecular features.

